# A Workflow for Spatial Transcriptomic Analysis from Intra-operative Human Skeletal Muscle Biopsies

**DOI:** 10.64898/2026.02.24.707605

**Authors:** Patricia S. Pirbhoy, Vikram Murugan, Michael Hicks, Oswald Steward, Ranjan Gupta

**Author notes:** Co-first authors (Patricia S. Pirbhoy, Vikram Murugan). Co-senior authors (Michael Hicks, Oswald Steward, Ranjan Gupta). Corresponding author: Oswald Steward, Ph.D. Department of Anatomy and Neurobiology, School of Medicine, 837 Health Sciences Rd., Irvine, CA 92697, USA.

## Abstract

**Introduction:** Successful reinnervation following peripheral nerve injury is highly variable, and the molecular programs underlying human muscle degeneration and recovery remain poorly defined. There is a critical need for high-resolution, spatially resolved gene expression data from human skeletal muscle obtained in clinically relevant settings. This study aimed to establish the feasibility of applying spatial transcriptomics to intra-operatively human muscle biopsies and to generate a framework for identifying gene expression signatures associated with reinnervation outcomes.

**Methods:** To validate the workflow, we collected biopsies intraoperatively from upper-extremity muscles during standard-of-care orthopaedic surgical procedures 5 months after traumatic brachial plexus injury. The flash-frozen biopsy was processed using the 10x Genomics Visium HD high-resolution platform. Quality metrics confirmed high RNA integrity and robust transcript detection at 8 µm resolution.

**Results:** Genes involved in neuromuscular junction formation, degeneration, and regeneration were identified at subcellular resolution and showed fiber-type-specific expression patterns. Analyses were performed using complementary approaches in Seurat and Loupe Browser.

**Conclusions:** Together, these findings demonstrate the feasibility of spatial transcriptomics in human muscle, establish baseline gene-expression signatures, and provide a foundation for future studies aimed at identifying biomarkers associated with successful reinnervation and improved nerve-repair strategies.

## Introduction

Muscle denervation following peripheral nerve injury produces immediate paralysis, progressive muscle atrophy, and functional decline. While nerve repair or transfer can re- establish axonal contact with the distal end of the transected nerve, functional recovery of meaningful strength remains inconsistent due to reinnervation failure into the recipient target muscle. Prior work from our team has used preoperative muscle biopsies to evaluate neuromuscular junction integrity and degeneration after denervation to identify factors contributing to reinnervation success vs. failure after nerve transfer (Chen et al., 2023). A key finding is the surprising, prolonged survival of motor-endplates (MEPs) in denervated muscle biopsies in some patients that exhibit successful reinnervation (Gonzales et al., 2025). These findings suggest the possibility that preservation of MEPs might predict reinnervation competence. However, assays of MEP morphometry require elaborate preparations and quantitative analysis and likely only provide a partial proxy for the broader transcriptional and microenvironmental changes that govern functional reinnervation.

Accordingly, we set out to explore whether a signature of reinnervation competence can be identified through high-resolution spatial transcriptomics using the 10x Genomics Visium HD spatial transcriptomic platform. Conventional single-cell RNA-seq requires tissue dissociation and thus loses spatial context; single-nucleus RNA-seq retains architecture but loses extranuclear RNA transcript information. High-definition (HD) spatial transcriptomics overcomes these limitations by unbiased quantification of transcripts in the spatial architecture of tissue. Visium® HD deploys a barcode-based technology that can capture gene expression profiles in 2, 8, or 16 µm^2^ bins, enabling transcriptomic mapping within sub-cellular domains of individual muscle fibers, structural matrix components, and their microenvironment.

Here, we describe a workflow and initial analysis of human muscle biopsies collected during routine orthopaedic procedures. Using a combination of Seurat v5 (R) (Hao et al., 2021, Satija, 2024, Hoffman and Satija, 2023) and Loupe Browser (10x Genomics), we evaluate quality control metrics of the spatial transcriptomic dataset, assess muscle fiber composition, perform cluster-based analysis to identify unique gene expression profiles, and investigate neuromuscular junction-associated genes that may contribute to successful innervation. Defining gene expression profiles of human muscle at different times post-denervation could identify biomarkers that predict successful reinnervation, providing patient-specific data to inform surgical decision-making and optimize timing of nerve repair.

## Materials and Methods

### Patient and Tissue Sample Information

A 24-year-old male patient undergoing standard-of-care orthopaedic surgery for traumatic brachial plexus injury was enrolled under an institutional review board (IRB)–approved protocol and provided informed consent. A trapezius muscle biopsy confirmed to be normally innervated preoperatively was obtained intraoperatively during left brachial plexus reconstruction involving cable sural nerve grafting to the musculocutaneous and median nerves five months after traumatic injury. Biopsies were flash-frozen in liquid nitrogen-cooled isopentane and stored at -80 °C before cryosectioning at 10 µm.

### Visium HD library preparation, sequencing, and image alignment

Sections were stained with hematoxylin and eosin (H&E) for histological quality assessment; one section per sample was selected and imaged at 10X magnification with a Nikon Ti-E widefield microscope (**Figure 1B**) for subsequent alignment with CytAssist images (Visium HD Fresh Frozen Tissue Preparation Handbook, Rev E; RRID: SCR_023571). The workflow for sample preparation is illustrated in **Figure 1A**. Other sections were taken for RNA Integrity Number (RIN); these were submitted separately to the UCI Genomics Research and Technology Hub (GRT Hub; RRID: SCR_026615) for RNA extraction. RNA concentration and purity were assessed using a Nanodrop spectrophotometer; RNA integrity was evaluated using the Agilent Bioanalyzer RNA pico assay.

**Figure 1.**
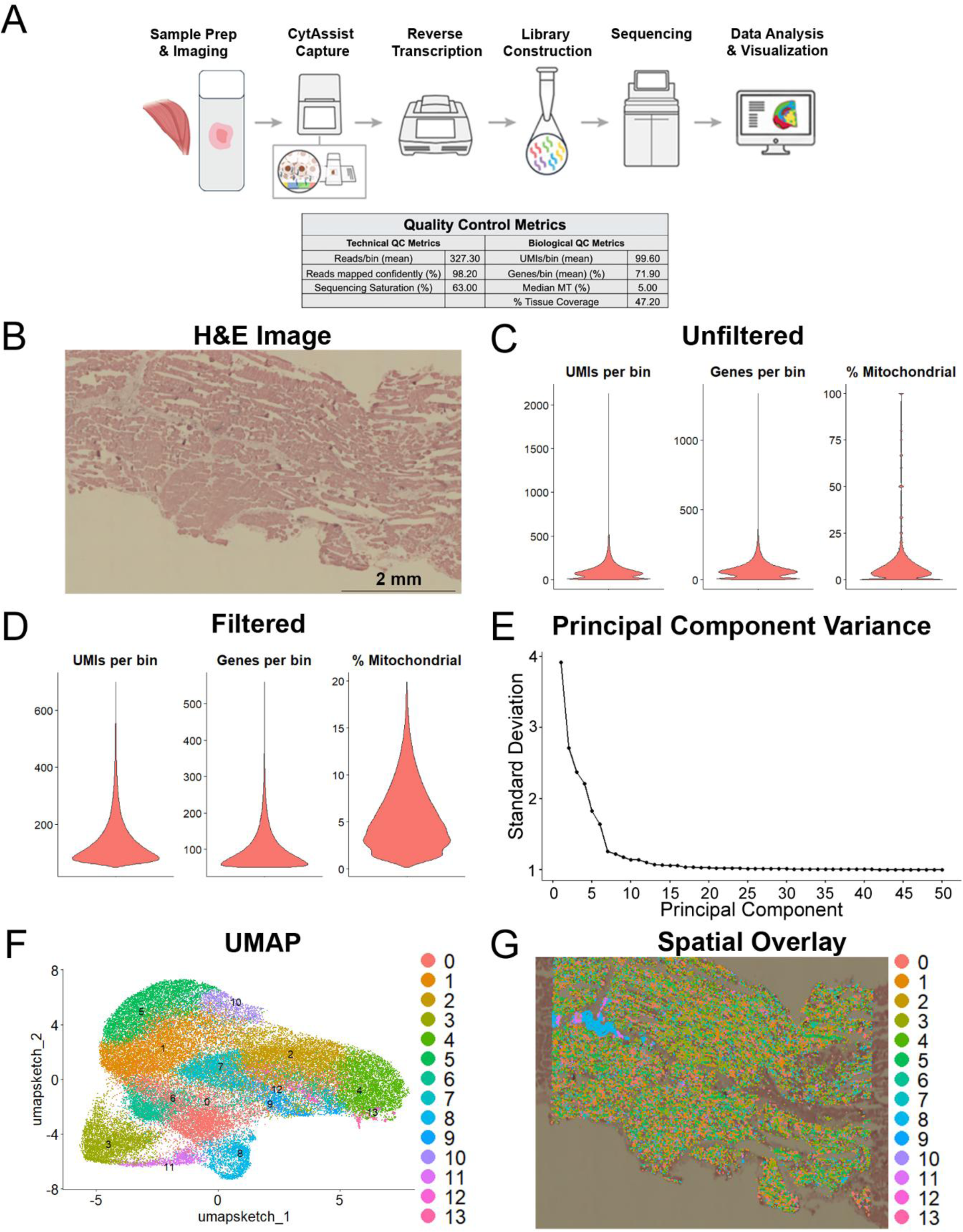
Quality control metrics for a muscle biopsy sample following spatial transcriptomics analysis. **A.** Overview of the Visium HD spatial transcriptomics workflow using the human whole-transcriptome panel (Image courtesy of 10x Genomics, Inc.). Table summarizes key technical and biological quality control metrics. **B.** H&E-stained human trapezius biopsy, scale bar is 0.5 mm. **C.** Violin plots of unfiltered data showing distribution of unique molecular identifiers (UMIs, total number of RNA transcripts detected) per 8 µm bin, genes per bin, and mitochondrial gene percentage. **D.** Violin plots of filtered data after applying thresholds for UMI (>20, <500), gene counts (>20, <300), and mitochondrial gene percentage (>0, <20%). **E.** Elbow plot showing the variance explained by principal components. **F.** Uniform Manifold Approximation and Projection (UMAP) visualization generated using the top 13 principal components (PCs) and a clustering resolution of 0.5. Parameter selection was guided by published Seurat and Visium HD vignettes. **G**. Seurat spatial overlay of transcriptional clusters mapped onto the aligned H&E image.

Selected sections were transferred to a 6.5 x 6.5 mm capture area on a Visium HD slide (10x Genomics, PN 1000675) via CytAssist-Enabled Probe Release & Capture. Gene expression libraries were generated from each tissue section and sequenced. The whole human transcriptome probe panel was used for these studies (Visium HD Spatial Gene Expression User guide, Rev D). Visium HD processing was completed by the UCI GRT Hub. Nikon Ti-E images, CytAssist images, and spatial transcriptomic outputs were then uploaded to the 10x Genomics Cloud Analysis for image alignment and generation of Loupe Browser v9.0.0 (RRID: SCR_018555) compatible file output (cloupe).

### Data processing and Seurat v5 analysis

Spatial transcriptomics data were processed using the Seurat package (RRID: SCR_016341) following the 8 µm Visium HD workflow described in the Satija Lab vignette (Hoffman and Satija, 2023, Satija, 2024). Violin plots were used to visualize the distribution of unique molecular identifiers (UMIs, total RNA transcripts per 8 µm region), genes detected per region, and mitochondrial gene percentage in the raw dataset (**Figure 1C**). Based on these distributions, thresholds were applied for UMI (>50, <700), gene counts (>50, <1000), and mitochondrial gene percentage (>0, <20%) to exclude low-quality or damaged tissue regions while retaining spots with sufficient transcript complexity. The filtered dataset was visualized using the same metrics (**Figure 1D)**. The elbow plot (**Figure 1E)** shows the variance explained by principal components (PCs). Dimensionality reduction was performed on the top 13 PCs using Uniform Manifold Approximation and Projection (UMAP), with a clustering resolution of 0.5, yielding 14 clusters (**Figure 1F).** The UMAP places bins with similar expressions near each other, creating clusters of similar data points.

Fiber-type-specific differential gene expression (DGE) analyses were performed in R (v4.3) using Seurat v5. The spatial transcriptomics object from the Visium HD dataset was subset to include only the innervated control sample. Composite marker scores were calculated for each spatial bin by averaging z-scored expression of canonical myofiber genes previously reported in the literature (Hsu et al., 2024). For the slow Type I and fast Type II comparison, marker sets included MYH7, TNNT1, ATP2A2 (Type I), and MYH1, MYH2, TNNT3, ATP2A1, TPM1 (Type II). For the fast-fiber subtypes, Type IIA fibers were defined by MYH2, TPM1, TNNT2, ATP2A1, and Type IIX fibers by MYH1, TPM1, TNNT3, and ATP2A1. Each spatial bin was assigned to the fiber type or subtype whose composite score exceeded the other by ≥0.10. Bins with ambiguous assignments were excluded.

DGE was performed within the innervated sample using the FindMarkers function from the Seurat package (McDavid et al., 2013). In the resulting output, positive log_2_ fold-change values indicate higher expression in Type I or Type IIA myofibers, whereas negative values indicate enrichment in Type II or Type IIX fibers, respectively. Results were exported as full and FDR < 0.05 CSV tables. All analyses were conducted on normalized “Spatial” assay data.

### Spatial autocorrelation and hotspot visualization

Spatial autocorrelation of gene expression was assessed using the global Moran’s I statistic implemented in the *spdep* R package (Bivand, 2022, Bivand and Wong, 2018). A predefined panel of NMJ-associated genes (AGRN, LRP4, CHRNG, CHRNA1, PREP, CHRND, CHRNB1, DOK7, VAMP1, MUSK, RAPSN, CHRNE, COL13A1) was selected, and global and local Moran’s I were computed for each gene individually to assess spatial clustering. Moran’s I measures the correlation between each bin’s expression value and those of its spatial neighbors to determine whether high or low expression values are spatially clustered. The statistic ranges from -1 (perfect spatial dispersion) to +1 (strong spatial clustering). A Z-score and corresponding p-value were computed under a randomization null model, with significant positive values indicating non-random spatial clustering.

To identify spatial hotspots, local indicators of spatial association (LISA) were computed for each gene, classifying bins as High-High (HH), Low-Low (LL), High-Low, or Low-High relative to neighboring expression values. Significant HH bins (p<0.05, after false discovery rate correction) were extracted and converted into spatial polygons to delineate discrete co-expression domains. To evaluate whether NMJ genes co-localized within the same spatial domains, HH assignments for the full gene panel were compiled into a spot-by-gene matrix and used to compute pairwise HH overlap across all gene combinations. Buffered HH maps were generated by expanding the perimeter of each significant HH cluster by one bin radius (∼8 µm) to include immediately adjacent regions of elevated expression. Buffered hotspots were then overlaid on the tissue image and visualized in red to highlight regions of enriched NMJ-associated gene expression.

## Results

### Intra-operative trapezius biopsy demonstrates high RNA quality for high-resolution spatial analysis

Muscle biopsies were collected in the operating room and immediately prepared for downstream processing using the 10x Genomics Visium HD workflow (**Figure 1A**). Pre-sequencing review of H&E-stained sections revealed no freeze artifacts or other defects (**Figure 1B**). Assessment of RNA Integrity Number (RIN) yielded 7.7, confirming intact RNA and overall sample integrity. Our initial evaluation of the spatial transcriptomics dataset focused on technical and biological quality metrics to ensure suitability for downstream analysis (**Figure 1A, bottom**). Technical quality control (QC) metrics indicated robust data capture and sequencing performance. On average, each 8 µm bin contained 327.3 reads, with 98.2% of reads confidently mapped and a sequencing saturation of 63%, consistent with adequate depth and library complexity for downstream spatial gene expression analysis. Each 8 µm bin contained 99.6 unique molecular identifiers (UMIs; total transcript counts), and 71.9 detected genes, with a median mitochondrial gene fraction of 5% and 47.2% tissue coverage. These values indicate well-preserved muscle tissue and sufficient RNA complexity for spatial transcriptomic analysis.

### Complementary pipelines, Seurat and Loupe Browser, reveal spatially coherent transcriptional domains

We next compared spatial gene expression patterns using Seurat and 10x Genomics Loupe Browser. Using the Seurat v5 workflow for analysis, visualization, and integration of Visium HD spatial datasets (Satija, 2024, Hao et al., 2024, Hie et al., 2019), we generated a Seurat object from 8 µm bins and applied standard log-normalization. An elbow plot generated from the full dataset guided principal component (PC) selection, identifying 13 PCs for downstream analysis (**Figure 1E**). Because full-resolution Visium HD data contain hundreds of thousands of spatial bins, direct UMAP computation is demanding. Accordingly, we used leverage-score sketching to create a representative subset of 5,000 bins that preserved global transcriptional diversity, including rare populations. The 3,000 most variable genes were identified and used for downstream dimensionality reduction and clustering. A clustering resolution of 0.5 yielded a two-dimensional UMAP comprising 14 clusters (**Figure 1F**). Each cluster represents spatial bins with similar gene expression profiles, and inter-cluster distances reflect relative transcriptional similarity.

Seurat visualizations lack the high-resolution H&E-alignment available in Loupe Browser, which limits direct anatomical correspondence (**Figure 1G**). In contrast, Loupe Browser integrates unsupervised clustering directly with tissue morphology and uses all spatial bins without sub-sampling. Raw Visium HD data from innervated trapezius were examined in Loupe Browser v8 using k-means (k=10), yielding ten baseline expression groups with a UMAP and matched spatial overlay (**Figure 2A**). Using the largest k-means value that generated 10 cluster populations on Loupe Browser provided a broad tissue domain prior to any filtering and served as a reference map for subsequent QC. Canonical marker genes were used to annotate clusters into categories, including fast and slow-twitch myofibers, as well as non-myofibers, including fibro-adipogenic progenitors (FAPs), immune, endothelial, peripheral nerve, and epithelial groups. Spatial localization was also referenced when determining cluster assignments. Loupe distribution plots were used to set barcode-level quality control filters. UMIs per barcode (Log2) were restricted to 4.05-9.20 (**Figure 2B**), and genes per barcode (Log2) to 4.17-8.37 (**Figure 2C**). Application of these thresholds removed 54,884 barcodes, of which 48,786 (89.6%) of the excluded barcodes localized to fatty/fibrotic regions on the tissue image (**Figure 2D & Figure 2A**), confirming that low UMI/gene signal predominantly arose from non-myofiber compartments. After setting QC metrics thresholds, re-clustering the filtered dataset in Loupe Browser produced 21 unbiased, higher-resolution, spatially coherent clusters.

**Figure 2.**
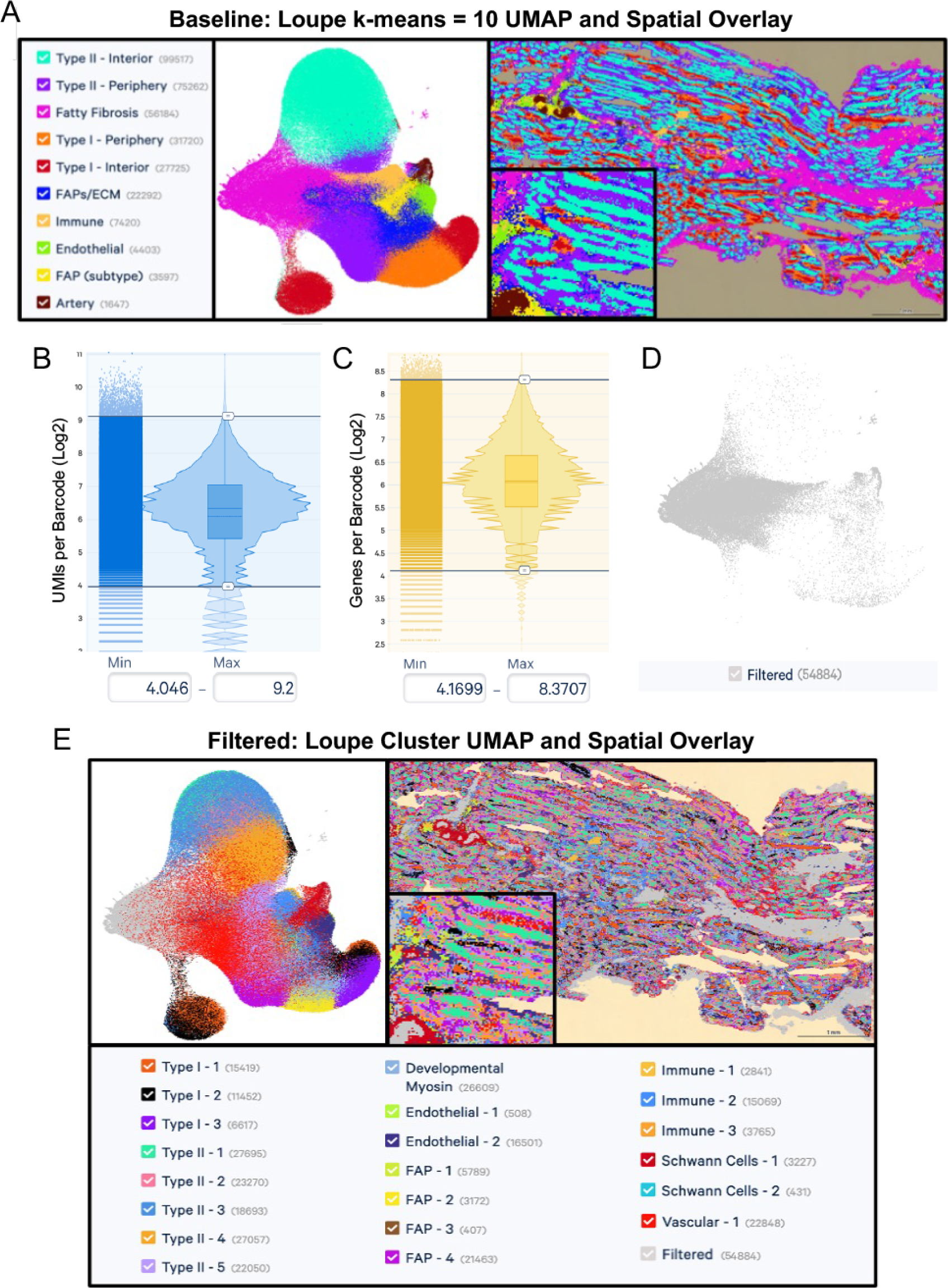
Loupe Browser clustering, QC filtering, and spatial mapping of trapezius Visium HD data. **A.** Unsupervised baseline clustering with Loupe k-means (k=10): UMAP of all barcodes colored by cluster (middle) with corresponding spatial overlay on the H&E image (right). Numeric labels corresponding to Loupe cluster IDs (left legend). **B.** Barcode-level QC: Log2 UMIs per barcode distribution used to set inclusion thresholds. **C.** Barcode level DC: Log2 genes per barcode distribution used to set inclusion thresholds. **D.** Barcodes removed by QC: UMAP highlighting the set excluded by the UMI and gene filters (low-information bins). **E.** Post-filter clustering: UMAP of retained barcodes colored by filtered Loupe k-means clusters (left), with the matched spatial overlay on H&E (right), and legend of cluster IDs (bottom).

These complementary approaches illustrate how Seurat’s computational framework enables efficient processing of large spatial datasets but lacks high-resolution visualization of gene expression profiles provided by Loupe Browser, which offers direct mapping of expression-defined domains within a histological context.

### Spatial transcriptomics captures cell-type heterogeneity and subcellular transcript localization within human myofibers

We next profiled the genes driving each cluster to define the major cellular populations present in the sample. Major population categories consisted of fast-fiber clusters (33.4%), slow-fiber clusters (12.2%), a developmental-myosin cluster (9.7%), fibro-adipogenic progenitors (11.2%), immune (7.9%), vascular (14.5%), and nerve (1.3%) groups (**Figure 2E**). Within several reported cluster categories (immune, FAPs, etc.), multiple genetically distinct clusters were identified, including four FAP clusters, three immune clusters, two endothelial clusters, and two nerve clusters.

In the myofiber compartments, multiple subclusters were detected that captured the layered architecture of individual myofibers, with clusters arising from the central contractile core separated from those localized to the sarcolemma rim. In the myofiber compartments, multiple subclusters were detected that correspond to different intracellular domains. A zoomed spatial view (**Figure 3A**) demonstrates Type I (slow) and Type II (fast) clusters that follow myofiber outlines and partition into domains within individual fibers, mapping a central contractile core (Orange: Type I; Green: Type II) and peripheral and sarcolemma domains (Black and Purple: Type I; Pink, Blue, Amber, Lavender: Type II). This spatial partitioning highlights how high-resolution spatial transcriptomics can capture not only major cell types but also <u>subcellular</u> <u>transcript localization</u> and structural heterogeneity within muscle fibers themselves. To illustrate the basis of our cluster labels, **Figure 3B** lists the top six differentially expressed genes (ranked by p-value) from the filtered Loupe Browser clusters shown in **Figures 2E** and **3A**. The ranked gene lists guided cluster annotations allowing us to link molecular identity with the unique spatial territories highlighted in **Figure 2E and 3A**. In practical terms, Type I fiber cores are enriched for slow-contractile genes (TNNC1, TNNT1, TNNI1), while the surrounding rim shows modulators of contraction and calcium cycling such as (MYLK3 and PPP1R1C). Type II fiber cores are concentrated with fast-contractile genes (MYH1, MYH2, MYBPC2) and calcium-handling markers (PVALB and CALML6). Beyond classification, this approach demonstrates the ability to visualize hundreds of transcripts, revealing their compartmentalization within intact muscle.

**Figure 3:**
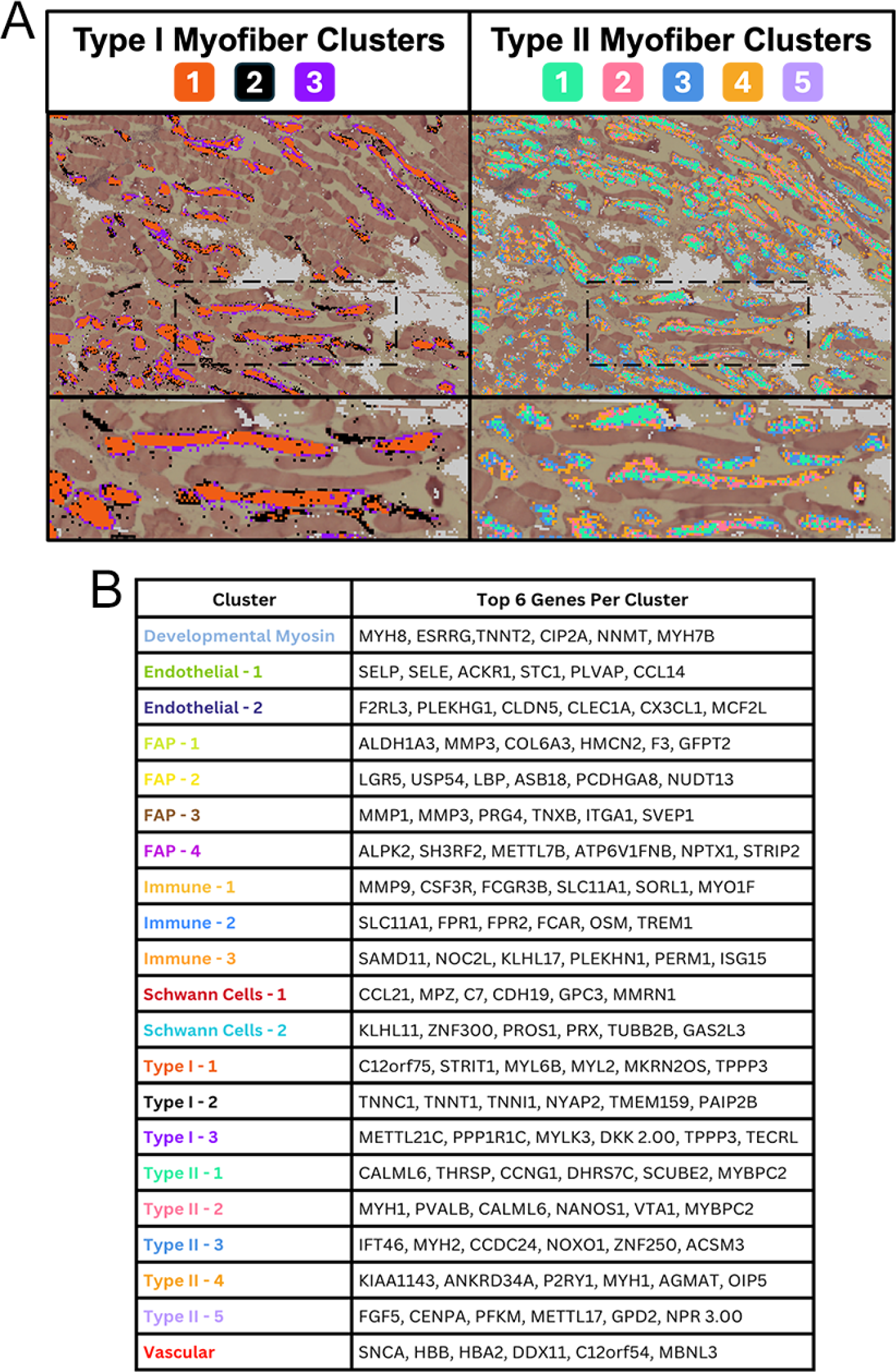
Fiber-type domains and representative genes in trapezius Visium HD Data. **A.** Zoomed spatial overlays of filtered Loupe Browser clusters highlighting Type I (slow) and Type II (fast) myofiber clusters, demonstrating layered myofiber domains. **B.** Representative markers for four clusters: Type I-1, Type II-1, Endothelial-2, and Vascular-1. Showing the top 10 genes per cluster ranked by differential-expression p value in Loupe Browser. Cluster colors and labels match those used in Figure 2E.

### Spatial transcriptomic mapping shows Trapezius Type II predominance and distinct Type IIA/IIX programs

To evaluate whether the spatial transcriptomic data accurately captured known fiber-type-specific gene expression profiles and to determine fiber type composition in the trapezius, we visualized canonical Type I and Type II myofiber markers using co-expression analysis in Loupe Browser (**Figure 4A-B)**. Since Loupe Browser performs differential gene expression analysis on selective clusters and not specific genes, we performed Seurat-based differential gene expression (DGE) analysis using a subsetted list of Type I (MYH7, TNNT1, ATP2A2) and Type II (MYH1/2, TPM1, TNNT3, ATP2A1) marker genes previously identified by Hsu et al., 2024 (**Figure 4A**, top table). The analysis was conducted using the Wilcoxon rank-sum test, with significance defined as FDR <0.05. In the Type I vs Type II DGE comparison (**Figure 4A**, bottom table), the top six upregulated genes for each fiber type are shown with their corresponding Log_2_ fold change (Log_2_FC) values.

**Figure 4.**
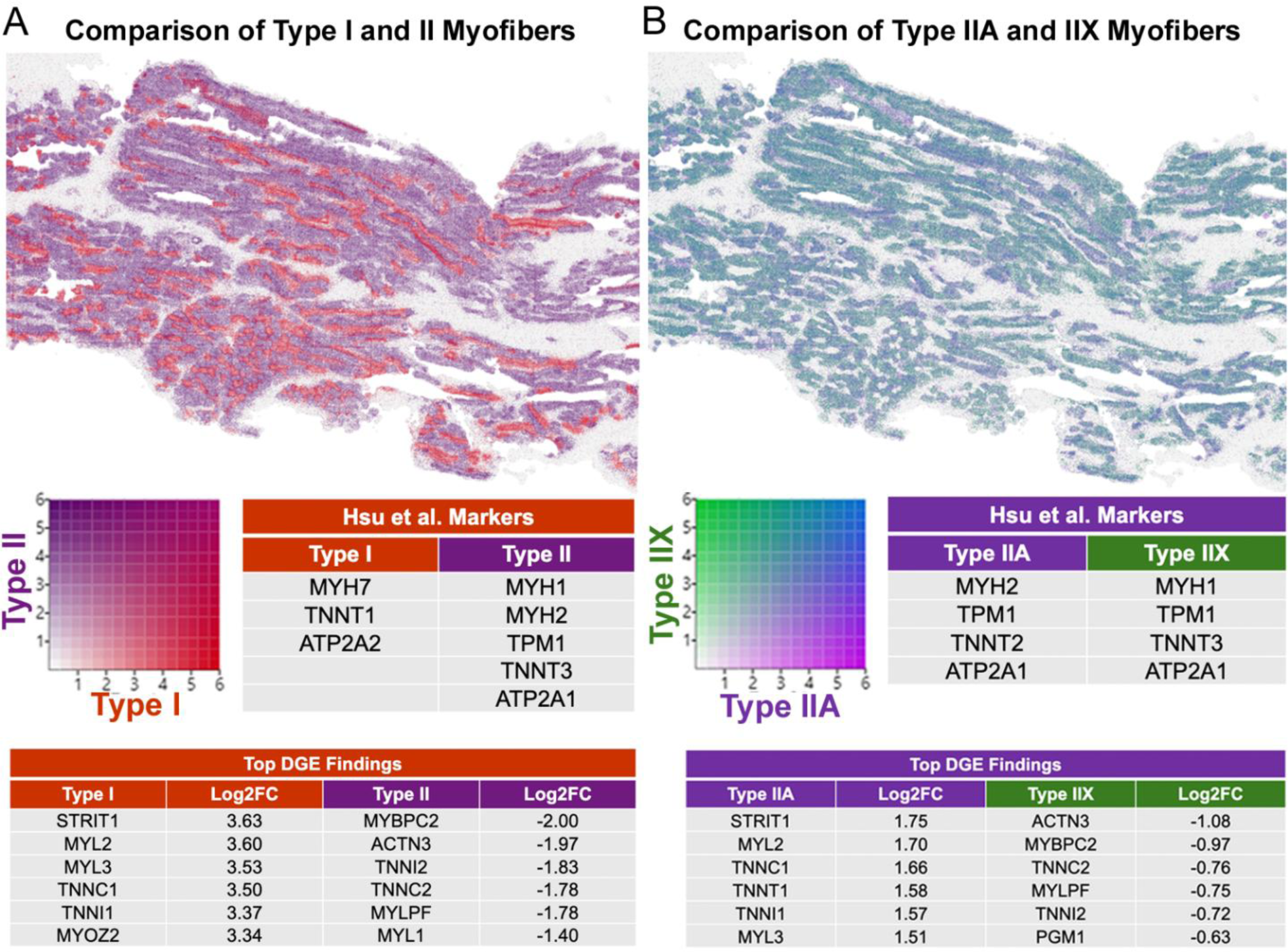
Differential gene expression analysis of human trapezius muscle using Visium HD spatial RNA sequencing. **A.** Loupe Browser co-expression map showing combined gene expression signatures for Type I and Type II myofiber groups in a human trapezius section, revealing a predominance of Type II-associated transcripts. Canonical marker genes from Hsu et al., 2024, were used to classify bins into Type I and Type II groups for downstream differential expression analysis (Table, top left). Differential gene expression (DGE) between Type I and Type II bins within the innervated control sample using the Wilcoxon rank-sum test (FDR < 0.05). Table shows the top six upregulated genes (ranked by log_2_ fold change, Log2FC) for each myofiber type following DGE analysis (bottom). **B.** Subclassification of Type IIA and Type IIX subtypes, using the corresponding marker genes from Hsu et al. 2024, identified spatial enrichment of Type IIX-associated transcripts within the same section. Tables display the subtype-specific marker sets (top right) and the top six upregulated genes for each myofiber subtype resulting from DGE analysis (bottom).

Genes upregulated in Type I myofibers showed the expected slow-twitch signature, with enrichment for sarcomeric and calcium-handling components characteristic of oxidative fibers. These included slow troponin subunits TNNI1 and TNNC1, slow myosin light chains MYL2 and MYL3, and structural regulators such as MYOZ2 and STRIT1, which support sustained contraction and calcium homeostasis. In contrast, Type II myofibers displayed the canonical fast-twitch transcriptional profile, marked by elevated expression of fast troponin subunits TNNI2 and TNNC2, fast myosin light chains MYL1 and MYLPF, and fast-fiber structural proteins MYBPC2 and ACTN3. Collectively, these genes represent well-established molecular hallmarks of slow- and fast-twitch muscle, confirming that the spatial data accurately resolve fiber-type-specific contractile molecular signatures.

We next compared Type IIA and Type IIX myofibers using the same marker-based strategy (**Figure 4B)**. Type IIA myofibers displayed the expected oxidative-glycolytic hybrid profile, with higher expression of genes associated with enhanced calcium handling and oxidative capacity, including STRIT1, MYL2, TNNC1, TNNT1, TNNI1, and MYL3. In contrast, Type IIX myofibers exhibited a fast, glycolytic signature, marked by elevated expression of ACTN3, MYBPC2, TNNC2, MYLPF, TNNI2, and the glycolytic enzyme PGM1, consistent with their role in rapid, high-force contractions. Notably, most subtype differences were modest in magnitude (Log2FC < 1), reflecting subtle yet coordinated isoform shifts that distinguish these closely related fast-twitch populations. Together, these results indicate that the trapezius sample is predominantly composed of Type II fibers, with a pronounced enrichment for Type IIA, while still capturing the canonical molecular distinctions across Type I, Type IIA, and Type IIX subtypes.

### Neuromuscular junction-associated transcripts form spatially distinct domains in human muscle

To connect transcriptional profiles with structural localization, we visualized canonical genes defining myofiber-type mosaicism and the spatial distribution of neuromuscular junctions (NMJs). **Figure 5A-B** shows the H&E-stained CytAssist image and higher-magnification view highlighting regions corresponding to Loupe Browser images in subsequent panels. **Panels C-P** display spatial feature plots (left) generated in Seurat and corresponding Loupe Browser zoomed-in images (right) showing spatial expression patterns for key fiber-type and NMJ-associated genes. MYH7 marks Type I (slow-twitch) fibers (**C-D**), MYH1 and MYH2 mark Type IIX and Type IIA (fast-twitch) fibers (**E-H**), and CHRNA1 (**I-J,** acetylcholine receptor alpha-1 subunit), MUSK (**K-L,** muscle-specific kinase), CHRNB1 (**M-N,** acetylcholine receptor beta-1 subunit), and LRP4 (**O-P,** low-density lipoprotein receptor-related protein 4) are enriched at NMJs. High-resolution visualization by Loupe Browser revealed focal clustering of NMJ-associated transcripts within muscle fibers, consistent with subsynaptic localization of AChR subunit transcripts.

**Figure 5.**
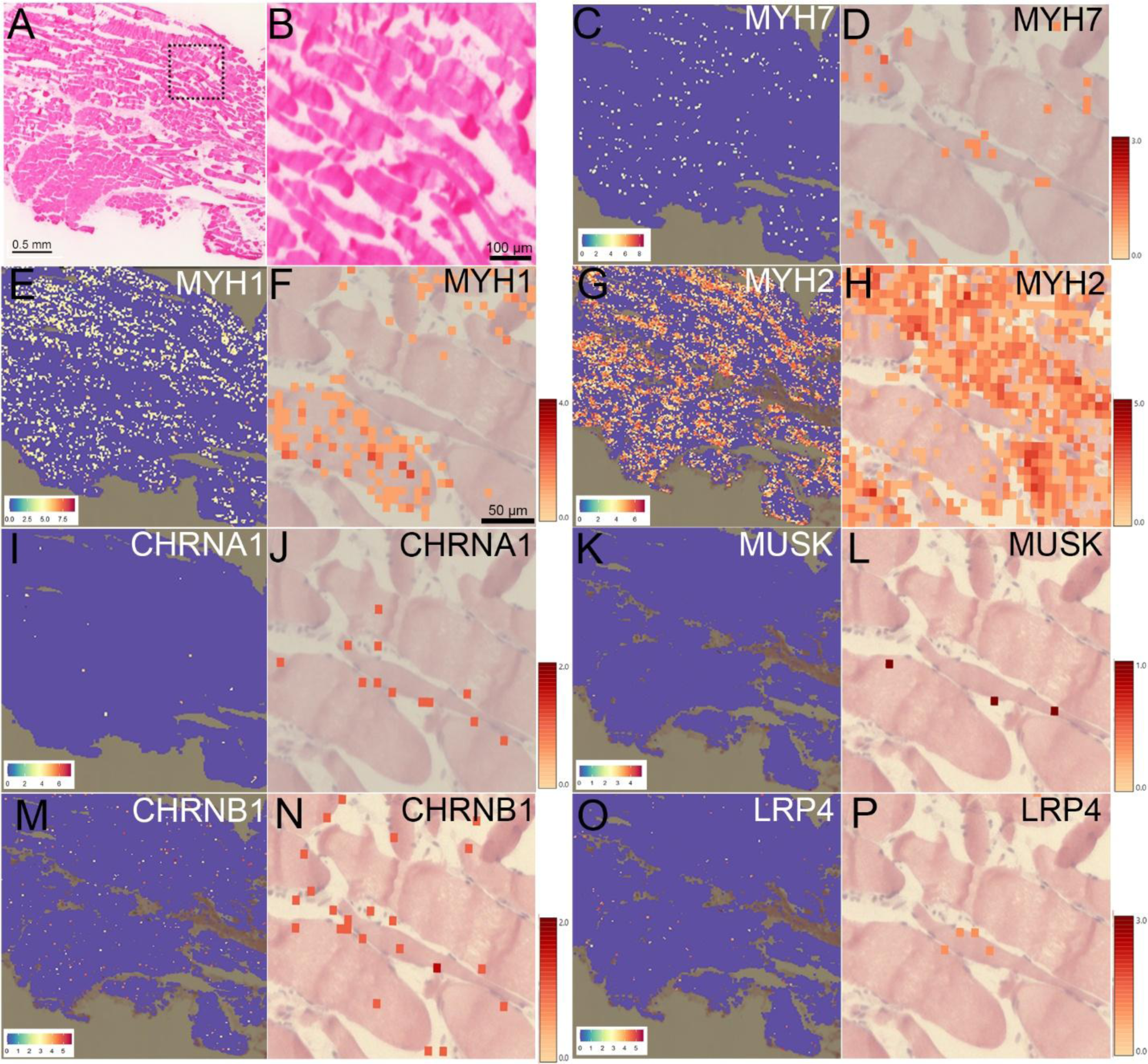
Fiber-type-specific and neuromuscular gene expression in innervated trapezius muscle. **A.** H&E-stained section imaged using CytAssist (Scale bar: 0.5 mm) **B.** High-magnification view of panel A showing the regions corresponding to Loupe Browser images in subsequent panels (Scale bar: 100 µm). **C-P.** Spatial Feature Plots (left) generated in Seurat and corresponding Loupe Browser zoomed-in images (right, scale bar: 50 µm) showing spatial expression patterns for key fiber-type and neuromuscular junction-associated genes. MYH7 marks Type I (slow-twitch) fibers (**C-D**), MYH1 and MYH2 mark Type IIX and Type IIA (fast-twitch) fibers (**E-H**), and CHRNA1 (**I-J,** acetylcholine receptor alpha-1 subunit), MUSK (**K-L,** muscle-specific kinase), CHRNB1 (**M-N,** acetylcholine receptor beta-1 subunit), and LRP4 (**O-P,** low-density lipoprotein receptor-related protein 4) are enriched at neuromuscular junctions. High-resolution imaging reveals focal clustering of CHRNA1 within muscle fibers, consistent with postsynaptic localization of acetylcholine receptor subunits. Heat map scales (right) indicate normalized expression intensity; note that ranges differ between genes.

To quantitatively assess whether the apparent clustering observed was statistically significant, we performed Moran’s I spatial autocorrelation analysis, which evaluates the degree to which gene expression values are spatially clustered or dispersed across tissue bins. Global Moran’s I analysis revealed significant positive spatial autocorrelation for several NMJ-associated genes, indicating non-random clustering within the tissue. Strong clustering was detected for CHRNA1, CHRNB1, CHRNG, CHRND, LRP4, AGRN, and PREP (Z-score > 10), and moderate but significant clustering for DOK7, VAMP1, MUSK, and RAPSN (Z-score = 2-10), whereas CHRNE (p=0.51) and COL13A1 (p=0.54) showed no significant spatial autocorrelation (**Table 1**).

**Table 1.**
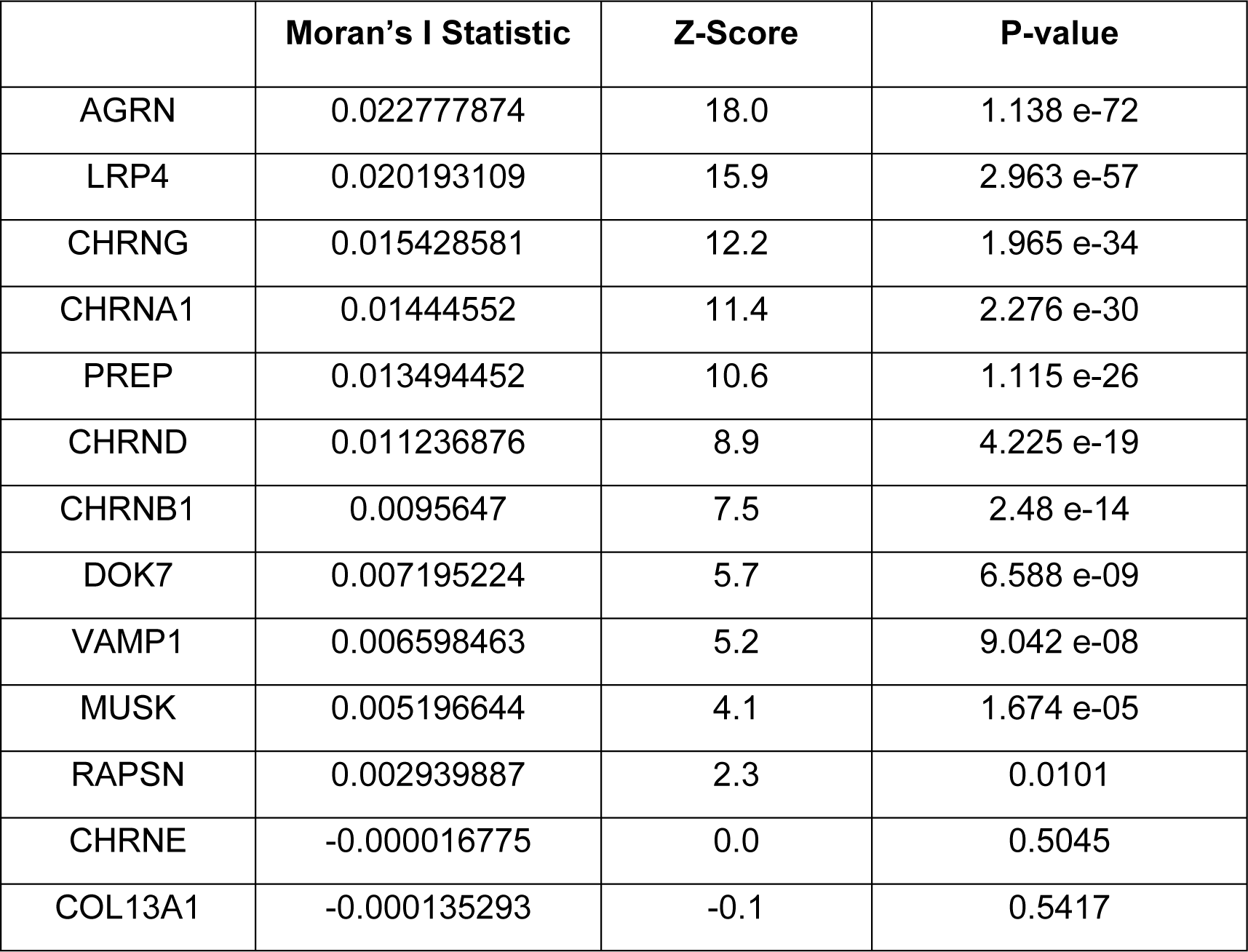
Spatial clustering of NMJ-associated genes in innervated trapezius muscle assessed by Moran’s I spatial autocorrelation. Spatial clustering of NMJ-associated genes was evaluated using Moran’s I spatial autocorrelation test. The table lists the Moran’s I statistic, Z-score, and p-value for each gene analyzed, reflecting the degree of non-random spatial distribution or dispersion within the tissue. Although Moran’s I values are numerically close to zero, consistent with subtle spatial effects at high resolution, positive Z-scores and low p-values indicate statistically significant clustering for most genes. In contrast, CHRNE and COL13A1 show no significant results, suggesting random spatial distribution. Moran’s I ranges from -1 (perfect spatial dispersion) to +1 (strong spatial clustering), with values near 0 indicating random spatial distribution.

To visualize these relationships locally, we performed a buffered High-High (HH) analysis, in which each bin was classified based on its expression value and the mean expression of its immediate neighbors (within a defined spatial radius). Bins exhibiting both high expression and high neighboring values were designated as HH regions and visualized in red (**Figure 6A**). These buffered HH maps highlight discrete hotspots of co-expression across the tissue section. By matching the location of these HH regions in Loupe Browser, we confirmed overlapping expression of CHRNB1 (**Figure 6B**) and LRP4 (**Figure 6C**) within the same localized domains (**Figure 6D**). Overall, these analyses demonstrate that NMJ-associated transcripts are not randomly distributed but are concentrated within discrete microdomains, reflecting spatially coordinated regulation of synaptic gene expression in muscle tissue.

**Figure 6.**
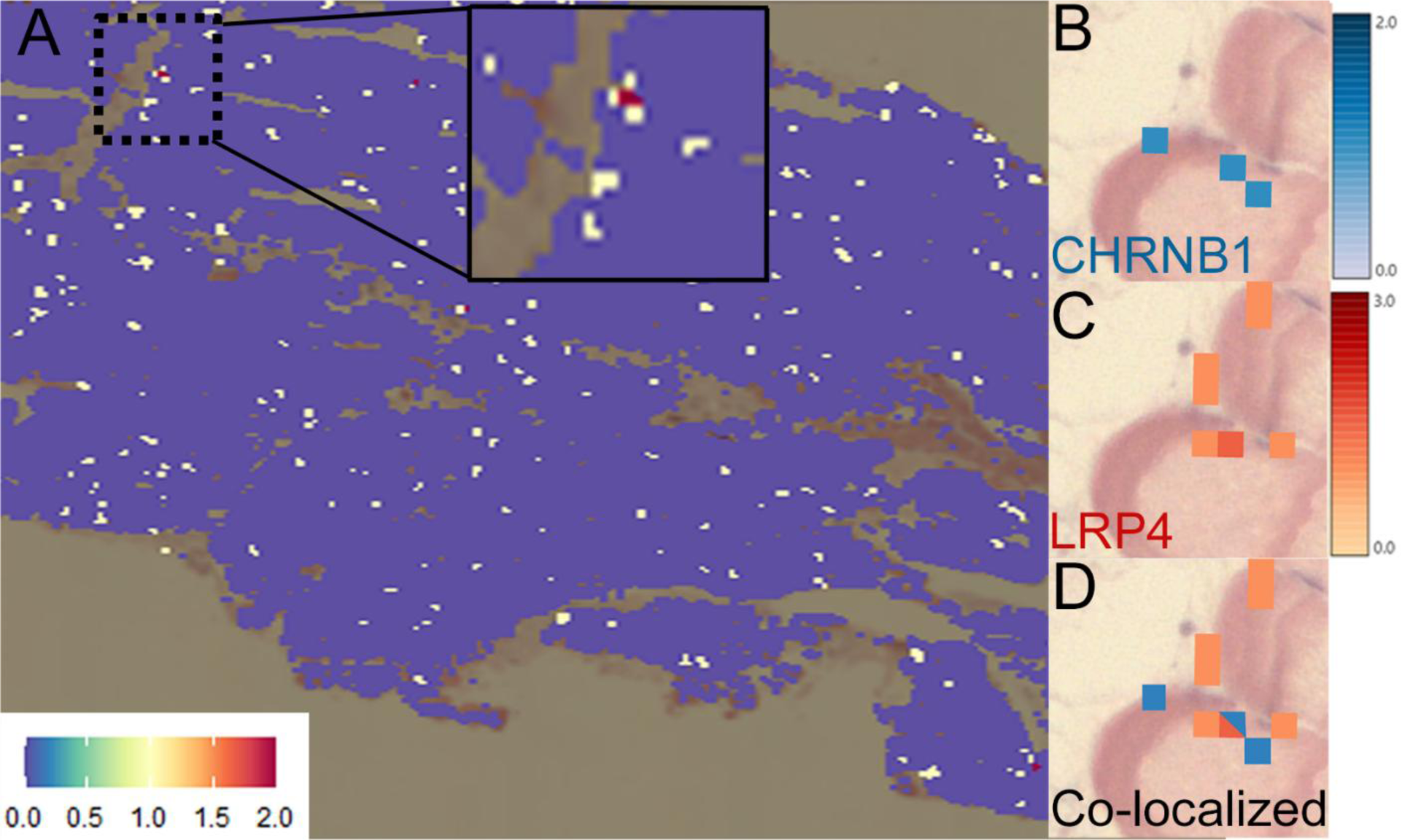
NMJ-associated transcripts are concentrated within discrete spatial domains. **A.** Buffered High-High (HH) expression map of NMJ-associated genes generated from Moran’s I indicators of spatial association (LISA) results. Regions of statistically significant co-expression (HH clusters) were buffered and visualized in red to highlight discrete transcriptional hotspots. Inset shows a zoomed-in region corresponding to the area visualized in panels B-D. **B.** Spatial localization of CHRNB1 expression visualized in Loupe Browser. **C.** Visualization of LRP4 expression. **D.** Colocalization of CHRNB1 and LRP4 within the identified hotspot.

Together, these results demonstrate that comparable spatial transcriptomic analyses can be performed using both Seurat and Loupe Browsers workflows. However, Loupe Browser’s integration of high-resolution imaging with tissue architecture enables precise subcellular localization of spatially distinct gene expression across diverse cell types and myofibers.

## Discussion

Our workflow demonstrates that high-definition spatial RNA-seq can be applied to human muscle biopsies obtained during routine orthopaedic surgeries. Using this approach, we mapped gene expression in sections of innervated trapezius muscle from a young male and resolved up to 22 transcriptionally distinct populations spanning myofibers and supporting cell types. Within Type I and Type II myofibers, we identified layered intracellular subdomains — core, peripheral, and sarcolemmal — each with unique gene expression profiles.. NMJ-associated transcripts, including CHRNA1, were sharply localized to discrete domains, enabling direct visualization of postsynaptic territories within individual muscle fibers. Together, these findings demonstrate that spatial transcriptomics can robustly characterize gene expression profiles of intra-operatively obtained human muscle, opening the door to studies of neuromuscular disease and injury. Importantly, the data generated can be made available broadly and analyzed using the programs of the platform (Visium) and other analysis programs.

### Advantages of Spatial Transcriptomics

Skeletal muscle is composed of different muscle fiber types and supporting cells. This presents a challenge for assessing gene expression with unbiased transcriptomic platforms that use tissue dissociation. The inability of dissociative techniques to resolve individual myofibers or their subcellular territories leaves key aspects of human muscle organization (myofiber types and subcellular domains within myofibers) unresolved. These features are critical for understanding the cellular and molecular changes that occur in neuromuscular disease or with denervation and reinnervation. To date, nearly all transcriptomic studies of human skeletal muscle have relied on dissociative approaches, yielding only coarse spatial information, limiting the interpretation of multinucleated fibers, myonuclear domains, and neuromuscular junction regions. To our knowledge, only a single study has applied spatial transcriptomics to human skeletal muscle biopsies, where the architecture necessary to understand the interplay between fibers and their surrounding niche is preserved (Umek et al., 2025). Umek et al demonstrated clear subcellular domains, with higher transcript density in nuclear and perinuclear zones, underscoring the importance of spatially resolved profiling for interrogating muscle architecture. A limitation, however, was that Umek et al used the Xenium platform with a predefined 377-gene panel and manual fiber segmentation to define transcript-density between fiber types and perimysium regions. In contrast, Visium HD provides whole-transcriptome profiling and derives unsupervised, expression-defined populations. Accordingly, this platform supports discovery-oriented mapping using native spatial context. Unsupervised clustering with immediate tissue localization reduces reliance on predefined regions and cluster annotation bias, which can help in detecting subcellular organization, such as myofiber core versus peripheral rim area (Min et al., 2025), and domains enriched in transcripts critical to the neuromuscular junction (more on this below). Such spatial precision is essential for understanding processes that depend on local mRNA translation, including at the sarcolemma and neuromuscular junction.

### Spatial Transcriptomics Identifies Subcellular Domains in Myofibers: Core vs. Peripheral Rim

Using Visium HD to profile a human trapezius biopsy at 8-µm resolution, we found that Type I (12.2%) and II (33.4%) myofibers together comprise 55% of the sectioned area, partitioning into intramyofiber domains corresponding to a central contractile core and peripheral layers. We also found tissue read depth reflected the spatial patterns that reflect underlying biology rather than under-sequencing, such as fatty-fibrotic areas, which contain little to no transcripts. Dissociative methods lose this within-fiber layering and concurrent fiber-type (Type I vs. Type II) spatial analysis, further underscoring the added value of spatial resolution.

Practically, our spatial readout enables comparison of gene programs at the core of the myofiber with those near the membrane. The CytAssist brightfield image and H&E alignment anchors transcript maps to intact anatomy. Recent high-resolution spatial approaches specifically underscore these advantages of spatial methods for muscle biology and disease mapping, where architecture and cell-cell relationships are readouts (Hoffman and Satija, 2023, Giordani et al., 2019). Furthermore, spatial profiling enables future multimodal integration, including alignment with imaging, electromyography, or histopathology, to connect molecular signatures with anatomical and functional readouts.

### Spatial Transcriptomics Defines Postsynaptic Microdomains

A key advantage of our spatial approach is the ability to localize transcripts within functional subcellular domains of human muscle and identify spatially coordinated clustering. By applying spatial autocorrelation and buffered High-High (HH) analysis, we found that multiple NMJ-associated transcripts were significantly clustered in discrete subcellular microdomains.

We first examined acetylcholine receptor (AChR) subunit genes (*CHRNA1, CHRNB1*, *CHRNG*); these are of particular interest because AChR clustering at the NMJ is key to neuromuscular transmission and because their localization depends on innervation (Hsu et al., 2024, Tsay and Schmidt, 1989). We then expanded to a panel of other genes previously identified as NMJ-associated genes. Spatial statistics revealed non-random, tightly localized expression patterns across the tissue for some of these other genes, demonstrating that NMJ transcripts other than AChR genes are highly localized in presumed postsynaptic domains. Notably, *AGRN* and *LRP4* – key organizers of AChR clustering – showed the strongest spatial clustering, as indicated by their high Moran’s I Z-scores, revealing a conserved molecular architecture consistent with their role as master regulators of synaptic organization. To our knowledge, this represents one of the first demonstrations of spatially co-localized NMJ transcript clusters in human skeletal muscle using an unbiased, whole-transcriptome approach. This level of subcellular resolution establishes a critical bridge between mechanistic insights from animal models and the molecular anatomy of the human neuromuscular junction. While model organisms have illuminated diverse muscle-resident populations and injury responses (Giordani et al., 2019), our findings provide direct evidence that synaptic microdomains can be mapped and molecularly characterized in samples obtained during routine surgeries.

### Human-Specific Insights from Whole-Transcriptome Spatial Profiling

High-resolution Seq-Scope was recently used to map normally innervated mouse soleus muscle and following denervation (3- and 7-days post). Through segmentation-based analysis, Type I, Type IIA, and Type IIX myofibers were resolved, highlighting fiber-type transcript programs, involving tropomyosins, troponins, and endoplasmic reticulum calcium pumps (Hsu et al., 2024 *Preprint*). Their marker sets aligned with prior snRNA-seq datasets from mouse soleus that distinguish myofiber types (Dos Santos et al., 2020, Petrany et al., 2020). Using the Hsu et al. gene list as an external reference for our dataset, our human trapezius analysis recapitulated contractile and calcium-handling signatures expected in Type I versus Type II and Type IIA versus Type IIX signatures. Notably, our whole transcriptome approach also revealed additional differentially expressed genes not reported in mouse studies. This difference could be due to the platforms used (Visium vs. Seq-Scope), muscle type (soleus vs. trapezius), or, more interestingly, to potential human-specific fiber-type programs or isoform shifts.

Complementing gene-oriented mapping, another study used a spatial-oriented approach with MALDI/AP-MALDI to draw metabolite-based maps that cleanly separated slow-oxidative from fast-glycolytic muscle territories, then confirmed them spatially with targeted RNA capture via LCM-RNA-seq (Luo et al., 2023). Notably, their spatial metabolomics also resolved a more oxidative-leaning subset within Type IIB fibers, illustrating that compartmentalization can exist within traditional fiber classes. Together, Hsu et al. and Luo et al. illustrate two complementary approaches to defining muscle fiber organization. Guided by these approaches, our human trapezius workflow integrates spatial segmentation (yielding Type I/II fiber distinction) and genetic anchoring, with top-gene signatures (Type I: TNNI1/TNNT1/MYL2/MYL3; Type II: MYH1/MYH2/ATP2A1/TNNI2) that extend these prior approaches to capture high-resolution myofiber subdomains. This joint framework produces a spatially and genetically resolved view of myofiber compartments, enabling finer stratification beyond slow/fast classes and setting up targeted myofiber and NMJ differential-expression analyses.

### Integrating Anatomical and Computational Frameworks for Spatial Analysis

For analysis, Loupe Browser provided a higher-resolution, anatomy-aware visualization framework than Seurat, enabling direct, high-magnification inspection of H&E-aligned tissue. This made intramyofiber vs. extramyofiber organization more readily apparent. Filtering for low UMI and gene counts in the trapezius removed regions of fatty or fibrotic tissue on the tissue image, indicating that low-UMI/low-gene bins reflect the low-cellularity fibrofatty areas and illustrating that RNA is unevenly distributed across the tissue. Re-clustering created Type I and Type II fiber subdomains and genetically distinct non-myofiber populations. Seurat complements Loupe Browser by enabling QC visualization, principal-component (PC) selection, and exploratory gene-level analysis. Using Seurat, we identified 13 PCs by elbow analysis and visualized bins with UMAP, which aided in exploratory comparison even when final per-section annotations were made in Loupe Browser (Satija, 2024, Coulis et al., 2023).

### Caveats and Limitations

Our study is limited by its single-subject, single-sample design. Our findings should be interpreted as an initial demonstration of feasibility rather than definitive conclusions about human muscle biology. Biological patterns we observed, including fiber-type programs and neuromuscular-junction transcript localization, are consistent with established muscle physiology, but comparisons across sex, age, and clinical situation is essential. An additional caveat is that the muscle studied here, identified as “innervated” by clinical assessment, was from a patient undergoing orthopaedic surgery for traumatic brachial plexus injury (specifically musculocutaneous and median nerve denervation).

Although Visium HD provides near-subcellular spatial resolution (8-µm bins), large multinucleated myofibers span many bins, and transcript localizations must be inferred from patterns observed within these bins rather than captured at actual intracellular boundaries. As a result, core vs peripheral rim domains represent spatially enriched and distinct zones rather than discrete structural compartments.

From a methodological perspective, only a subset of the sampled trapezius was profiled using Visium HD, as the workflow required trimming the biopsy to allow positioning on the Visium HD slide. This selection may not capture spatial heterogeneity within or across the sampled muscle. Looking ahead, incorporation of multiple denervated and normally innervated biopsies will require analytical frameworks capable of resolving shared versus subject-specific transcriptional states. Integrative tools, including Harmony (R) and scVI (Python), offer suitable approaches for multi-sample alignment, batch correction, and comparative analyses (Hao et al., 2021, Gayoso et al., 2022). 10x Cloud Analysis provides sample-integration workflows for standard Visium gene-expression data, provided the inputs meet size and format requirements; however, integration of Visium HD multiple datasets is not currently supported. As a result, cross-sample merging and integration must be performed in R or Python to enable robust comparative analyses.

Finally, although spatial transcriptomics is transformative for discovery, it is not feasible to deploy it routinely across large patient cohorts due to cost, tissue-handling requirements, and processing time. Our long-term objective is to use spatial transcriptomics as a discovery platform to define spatially anchored biomarkers of muscle fiber type, denervation, and reinnervation potential. Once identified and validated, such biomarkers may be translated into more scalable assays that are practical for broader clinical applications.

### Implications for Reinnervation Biology and Clinical Decision-Making

Our study demonstrated that muscle biopsies obtained during routine surgical procedures can be profiled at near-subcellular resolution, capturing features that are lost using traditional cross-sections or dissociative approaches. This workflow provides the spatial framework needed to assess these NMJ- and fiber-type signatures to denervated muscle. Because nerve-repair strategies and muscle-targeted therapeutics depend on the timing and quality of reinnervation, these readouts offer a path toward identifying biomarkers of successful reinnervation to inform surgical decision-making for nerve transfers at delayed post-injury intervals.

Beyond surgical applications, spatial transcriptomics provides a means for unbiased interrogation of muscle gene expression and transcript localization with disease and in response to therapeutic candidates. For example, mapping drug-responsive programs at cellular and subcellular levels can assist in optimizing therapeutic windows and prioritizing candidate targets for NMJ preservation in diseases involving the neuromuscular junction and with injury. Furthermore, our workflow can be extended to denervated cohorts and integrated with electrophysiology and clinical outcomes to prioritize candidate targets for NMJ preservation and translate spatial biology into actionable care for neuromuscular recovery.

## Author Contributions

P.S.P. and V.M. wrote the initial draft, generated figures and visualizations, and performed data analysis under the guidance of M.H. R.G. collected muscle biopsies during standard of care orthopaedic surgeries under an approved IRB (#20152278). All authors contributed to study design, data interpretation, conceptualization, and manuscript editing.

## Abbreviations

AChR: acetylcholine receptor
ACTN3: actinin 3
AGRN: agrin
ATP2A1: ATPase sacroplasmic/endoplasmic reticulum calcium transporting 1
CALML6: calmodulin-like 6
CHRNA1: cholinergic receptor, nicotinic, alpha 1
CHRNB1: cholinergic receptor, nicotinic, beta 1
CHRND: cholinergic receptor, nicotinic, delta
CHRNE: cholinergic receptor, nicotinic, epsilon
CHRNG: cholinergic receptor, nicotinic, gamma
COL13A1: collagen type XIII, alpha 1
DGE: differential gene expression
DOK7: docking protein 7
FAP: fibro-adipogenic progenitors
HD: high definition
HH: high-high
H&E: hematoxylin and eosin
IRB: institutional review board
LISA: local indicators of spatial association
LCM: laser capture microdissection
LRP4: low density lipoprotein receptor-related protein 4
AP-MALDI: atmospheric pressure matrix-assisted laser desorption/ionization
MEP: motor-endplate
MUSK: muscle-specific kinase
MYBPC2: myosin binding protein C
MYH1: myosin heavy chain 1
MYL: myosin light chain
MYLK: myosin light chain kinase
MYOZ2: myozenin 2
PC: principal components
PREP: prolyl endopeptidase
PVALB: parvalbumin
RAPSN: receptor-associated protein of the synapse
RIN: RNA integrity number
RNA: ribonucleic acid
RNA-seq: ribonucleic acid sequencing
snRNA: single nucleus
STRIT1: small transmembrane regulator of ion transport 1
TNNI1: troponin I type 1
TNNT1: troponin T type 1 TPM1 tropomyosin 1
UMAP: uniform manifold approximation and projection
UMI: unique molecular identifier
VAMP1: vesicle-associated membrane proteins

## Acknowledgments

Spatial transcriptomic kits were funded by NIA R21AG078909 (M.H. and R.G.). This work utilized resources of the UCI Genomics Research and Technology Hub (GRT Hub), parts of which are supported by NIH grants to the Comprehensive Cancer Center (P30CA-062203) and the UCI Skin Biology Resource Based Center (P30AR075047) at the University of California, Irvine, as well as to the GRT Hub for instrumentation (1S10OD010794-01and 1S10OD021718-01). P.S.P. was supported by the FRAXA Research Foundation Fellowship. Image detailing methods workflow in Figure 1A courtesy of 10x Genomics, Inc. A special thank you to Kelly Matsudaira for technical assistance and guidance with histological techniques.

## Ethical Publication Statement

We confirm that we have read the Journal’s position on issues involved in ethical publication and affirm that this report is consistent with those guidelines.

## Disclosures

O.S. is a co-founder and has economic interests in the company Axonis Inc., which is developing novel therapies for neurological disorders. All other authors have no disclosures to report.

## Disclaimers

None

## Statement of Clinical Relevance

This study demonstrates the feasibility of applying high-resolution spatial transcriptomics to human muscle biopsies obtained during surgery. Mapping fiber-type-specific and neuromuscular junction gene expression provides a framework for identifying molecular signatures associated with muscle degeneration, recovery, and reinnervation after nerve injury, laying the groundwork for future diagnostic and therapeutic studies in patients.

